# Developmental regulation of zinc homeostasis in differentiating oligodendrocytes

**DOI:** 10.1101/2023.07.26.550230

**Authors:** Christopher M. Elitt, Madeline M. Ross, Jianlin Wang, Christoph J. Fahrni, Paul A. Rosenberg

**Author notes:** Corresponding author: Christopher M. Elitt, MD, PhD Department of Neurology Boston Children’s Hospital 300 Longwood Avenue Boston, MA 02115. co-first authors (Christopher M. Elitt, Madeline M. Ross). co-senior authros (Christoph J. Fahrni, Paul A. Rosenberg).

## Abstract

Oligodendrocytes develop through well characterized stages and understanding pathways regulating their differentiation remains an active area of investigation. Zinc is required for the function of many enzymes, proteins and transcription factors, including those important in myelination and mitosis. Our previous studies using the ratiometric zinc sensor chromis-1 demonstrated a reduction in intracellular free zinc concentrations in mature oligodendrocytes compared with earlier stages (Bourassa et al., 2018). We performed a more detailed developmental study to better understand the temporal course of zinc homeostasis across the oligodendrocyte lineage. Using chromis-1, we found a transient increase in free zinc after developing oligodendrocytes were switched into differentiation medium. To gather other evidence for dynamic regulation of free zinc during oligodendrocyte development, qPCR was used to evaluate mRNA expression of the major zinc storage proteins metallothioneins (MTs), and metal regulatory transcription factor 1 (MTF-1) which controls expression of MTs. MT-1, MT-2 and MTF1 mRNAs were all increased several fold in mature oligodendrocytes compared to developing oligodendrocytes. To assess the depth of the zinc buffer, we assayed zinc release from intracellular stores using the oxidizing thiol reagent 2,2’-dithiodipyridine (DTDP). Exposure to DTDP resulted in a ∼100% increase in free zinc in developing oligodendrocytes but, paradoxically more modest ∼60% increase in mature oligodendrocytes despite the increased expression of MTs. These results suggest that zinc homeostasis is regulated during oligodendrocyte development, that oligodendrocytes are a useful model for studying zinc homeostasis in the central nervous system, and that regulation of zinc homeostasis may be important in oligodendrocyte differentiation.

## 1. Introduction

Oligodendrocytes are necessary for efficient propagation of action potentials in the brain and are increasingly recognized to serve dynamic roles in neuronal signaling, learning, and metabolism (Bonetto et al., 2020; Xin and Chan, 2020). During development, oligodendrocytes progress through a well characterized lineage prior to producing myelin (Kinney and Volpe, 2017). Aberrant white matter development leads to pathologies such as leukodystrophies and white matter injury of prematurity (Bugiani et al., 2013; Back and Rosenberg, 2014; Elitt and Rosenberg, 2014; Volpe, 2019; Garcia et al., 2020). In adults, endogenous oligodendrocyte progenitors can be mobilized for myelin repair following injury, such as in multiple sclerosis (Chang et al., 2002; Back et al., 2005), white matter stroke (Sozmen et al., 2016; Sozmen et al., 2019) and trauma (Rodriguez et al., 2014; Mendonca et al., 2021). A common theme in these pediatric and adult white matter disorders is oligodendrocyte differentiation arrest (Back et al., 2005; Nakahara et al., 2009; Bugiani et al., 2013; Cree et al., 2018; Srivastava et al., 2020). The ability to therapeutically promote differentiation of oligodendrocytes during these disease states requires a detailed understanding of cellular regulators of oligodendrocyte differentiation.

There has been considerable investigation of molecular mechanisms of oligodendrocyte differentiation (Adams et al., 2021; Clayton and Tesar, 2021), but these have generally neglected the role of brain micronutrients in oligodendrocyte development. This gap in understanding is in part due to a lack of appropriate tools to measure and visualize micronutrients in oligodendrocytes and other cells. Zinc is a micronutrient obtained exclusively through diet that was initially recognized as critical for human brain development in the early 1960’s after zinc-deficient children were noted to have growth and cognitive delays (Prasad et al., 1961). Zinc is the second most abundant micronutrient in the brain and is now recognized to serve diverse roles during development, injury and neurological disease (Sensi et al., 2009; Elitt et al., 2019).

Zinc is found in both bound and unbound forms in cells. While total zinc in the brain is abundant (∼100-300 μM), free zinc is kept relatively low (low nanomolar to picomolar range) (Maret, 2017). This dichotomy is because the majority of zinc is bound to proteins; indeed zinc binding proteins constitute 10% of the proteome (Andreini et al., 2006). A large portion of intracellular zinc is bound to metallothionenins or transcription factors, or sequestered in intracellular compartments such as mitochondria, lysosomes, nuclei, zincosomes or synaptic vesicles (Frederickson et al., 2005; Abiria et al., 2017; Maret, 2017). Because zinc plays an important role and excess free zinc can be toxic including to oligodendrocytes (Zhang et al., 2006; Domercq et al., 2013; Mato et al., 2013), there is tight homeostatic control of zinc in the brain (Frederickson et al., 2005; Kochanczyk et al., 2015) through a balance of the actions of zinc influx/efflux transporters and zinc storage proteins. The concentration of free zinc is in constant flux governed by the affinities of different proteins for zinc, changes in membrane permeability and release from intracellular compartments.

A key controller of zinc homeostasis is metal-responsive transcription factor-1 (MTF-1), which is a zinc finger transcription factor and the only identified metal-responsive transcription factor in mammals (Fukada et al., 2011; Gunther et al., 2012). Upon zinc binding, MTF-1 translocates to the nucleus where it binds to metal response elements within promoters for metallothioneins and the zinc efflux transporter ZnT1.

Zinc binding to MTF-1 directly couples changes in cytoplasmic zinc concentrations with transcriptional regulation of metallothionein and zinc transporter genes.

Characterizing how free zinc concentrations change across development and identifying regulators of zinc homeostasis in healthy developing oligodendrocytes is essential for understanding how zinc dyshomeostasis might contribute to oligodendrocyte differentiation arrest. Zinc dyshomeostasis is recognized to be involved in the pathogenesis of neurological disorders (Sensi et al., 2011; Scarr et al., 2016), including those involving white matter such as multiple sclerosis (MS) (reviewed in (Bredholt and Frederiksen, 2016)). Our prior work has demonstrated a decrease in free zinc concentrations in mature oligodendrocytes (Bourassa et al., 2018), suggesting that changes in zinc homeostasis may be involved in oligodendrocyte differentiation. In the present study, we sought to characterize changes in zinc across oligodendrocyte development in vitro, and to determine how these changes are accompanied by altered expression of zinc responsive proteins.

## 2. Materials and Methods

### 2.1 Oligodendrocyte Cultures

Stage-specific, highly enriched oligodendrocyte cultures were prepared from P2 rat forebrains modifying methods developed by McCarthy and colleagues (McCarthy and de Vellis, 1980) and previously described in detail from our laboratory (Rosenberg et al., 2003; DeSilva et al., 2009). P2 Sprague Dawley rat pups (Charles River, Wilmington, MA) were sacrificed by decapitation using sharp scissors without anesthetic.

Decapitation induces a rapid loss of consciousness and is an approved method of euthanasia by the American Veterinary Medical Association that minimizes rodent distress at this postnatal age. All procedures were also approved by the Boston Children’s Hospital Institutional Animal Care and Use Committee (IACUC) under protocols 20-03-4132R and 19-05-3901R and performed in accordance with NIH guidelines. Each culture used 9-12 mixed gender pups which were pooled.

Approximately 150 pups from 15 dams were used for these studies. Following forebrain dissociation, mixed glia cultures were grown in flasks for 10-17 days in DMEM with 10% fetal bovine serum in an incubator at 37°C with 5% CO2. Selective detachment was then used to isolate oligodendrocytes. Microglia were first removed using an orbital shaker set at 200 rpm for 1 hour, followed by a medium change. Shaking was continued for another 18 hours to remove oligodendrocytes, with astrocytes remaining adherent to the flasks. Oligodendrocytes were plated at a density of 10,000 cells/cm^2^ onto poly-DL-ornithine coated glass coverslips, 24 well plates, or onto 10 cm dishes in basal defined medium (BDM) supplemented with basic fibroblast growth factor (bFGF) (10ng/mL) and human platelet derived growth factor (PDGF) (10ng/mL) to promote proliferation. After 6 days, oligodendrocytes were transferred into differentiation medium (BDM) supplemented with triiodothyronine (T3) (3ng/mL) and ciliary neurotrophic factor (CNTF) (10ng/mL). Previous work has demonstrated that the majority of cells are at the oligodendrocyte progenitor stage after 2 days in proliferation medium (PM) containing PDGF and FGF, at the pre-oligodendrocyte stage after 6 days in PDGF and FGF and at the mature oligodendrocyte stage by 13-16 days following transfer into differentiation medium (DM) containing CNTF and T3 (Back et al., 1998).

Cultures are named by their eventual final stage: pre-oligodendrocyte (O4) in proliferation medium; mature oligodendrocyte (MBP) in differentiation medium.

### 2.2 Chromis-1 imaging

Chromis-1 is a novel ratiometric zinc sensor with numerous advantages over non-ratiometric zinc sensors, including the ability to detect physiologically relevant zinc concentrations (Bourassa et al., 2018). Methods for oligodendrocyte imaging are as previously described (Bourassa et al., 2018). Briefly, oligodendrocytes grown on 12 mm glass coverslips were incubated with chromis-1 (10μM) for 30 minutes at 37°C in BDM media without growth factors. Cells were then washed with Hanks Buffered Saline Solution (HBSS) containing calcium and magnesium. Coverslips were transferred to 60 mm dishes containing 4 mL HBSS and then imaging was performed using a LSM710 confocal microscope (Zeiss) equipped with a 2-photon laser (Spectra-Physics).

Excitation was at 720 nm and integrated emission intensities were collected through 2 band pass filters (425-462 nm) and (478-540 nm), corresponding to unbound and bound forms of the probe. Four unique fields of view were captured from each coverslip. An intensity ratio image depicting BP2/BP1 for each pixel was generated by Zeiss image analysis software and exported as 32-bit tiff files. These images were converted to 16- bit color images in ImageJ for subsequent analysis. An average ratio value was recorded for each cell using a region of interest (ROI) 30 pixels in diameter in the cytoplasm.

Experiments recording chromis-1 ratios in response to DTDP stimulation were performed using oligodendrocyte cultures prepared as described above. 100x stock solutions of 2,2’-dithiodipyridine (DTDP) and N,N,N’,N’-tetrakis(2-pyridinylmethyl)- 1,2-ethanediamine (TPEN) were prepared in dimethyl sulfoxide (DMSO). All chemicals were purchased from Sigma-Aldrich. DTDP and TPEN were directly pipetted onto coverslips to a final concentration of 100 µM and 3 µM, respectively. One image was obtained immediately before DTDP treatment, and then each minute after treatment for five minutes. Following five minutes of DTDP exposure, TPEN stock was pipetted onto cells, and an additional image was obtained after one minute. Image analysis was performed in ImageJ as described above. ROI x-y coordinates in ImageJ were used to ensure tracking of cells across the course of each experiment.

### 2.3 Quantitative real-time PCR

Primers:

Cyclin A (CyCA)

FWD: TATCTGCACTGCCAAGACTGAGTG REV: CTTCTTGCTGGTCTTGCCATTCC

MT1

FWD: GTTCGTCACTTCAGGCACAG REV: CGTTGCTCCAGATTCACCAG

MT2

FWD: CAGCTGCACTTGTCCGAAGC REV: GGACCCCAACTGCTCCTGTG

MT3

FWD: GACCTGCCCCTGTCCTACTG REV: CTGCATTTCTCGGCCTTGGC

MTF1

FWD: CCCCAACTCCTAACACGGCA REV: GCTACTGGTACTGCGGTGGT

Myelin Basic Protein (MBP)

FWD: CCAAGGAAAGGGGAGAGGCC REV: TTTTGGAAAGCGTGCCCTGG

Oligodendrocyte cultures were grown as above and harvested using the E.Z.N.A Total RNA Kit (Omega Bio-tek, Inc). For each 10 cm dish, 750 μL of TRK lysis buffer was added and cells lysed using a rubber cell scraper. The lysate was loaded onto a homogenizer mini column (Omega Bio-tek, Inc) and centrifuged to collect homogenized lysate. 750uL of 70% ethanol was added and samples vortexed. Samples were then loaded onto a HiBind RNA Mini Column and centrifuged at 10,000 x gravity for 1 minute. DNA was digested using RNase-Free DNase (Omega Bio-tek, Inc). Following 15- minute digestion, RNA was washed using RNA Wash Buffer I followed by 1 minute centrifugation. Steps were repeated with RNA Wash Buffer II diluted with 100% ethanol. RNA was eluted using 40 μL of nuclease-free water, centrifuged and then stored at -70°C. RNA concentration was determined and quality confirmed using an Agilent Bioanalyzer. cDNA was prepared with the Invitrogen High Capacity Reverse cDNA kit (ThermoFisher Scientific) using 1 μg of starting RNA with 10X RT Buffer (2.0μL), 25X dNTP Mix (100mM, 0.8μL), 10X RT Random Primers (2.0μL), MultiScribe Reverse Transcriptase (1.0μL) per 20μL reaction followed by PCR (c1000 Touch, Bio- Rad, Hercules, CA) for 10 minutes at 25°C, 120 minutes at 37°C, 5 minutes at 85°C, and then held at 4°C. Samples were then diluted with water to 140μL. For quantitative RT-PCR, cDNA (4μL) was added to a 96-well qPCR plate containing primers (0.5 μL of each primer, 500nM), water (5.5 μL) and 2x SYBR green (10μL, QIAGEN, Cat #330520) per 20μL reaction. The plate was sealed with adhesive film and qPCR was performed in a light cycler (Quant Studio 3, ThermoFisher Scientific) using the follow sequence: step 1: 95°C 10 minutes; step 2: 95°C 15 seconds; step 3: 60°C 1 minute; step 4: repeat steps 2-3 for 40 cycles. Routine control experiments to assess for contamination included no template and no reverse transcriptase controls. For a subset of the O4 DIV6 (n=3) and the MBP DIV16 (n=3) cultures, a similar procedure was carried out on a 384-well qPCR plate using identical primers and procedures.

Transcripts were normalized to the housekeeping gene CycA which was stable across oligodendrocyte development in these experiments and previously validated (Nelissen et al., 2010). Fold changes were calculated using the ΔΔCt method (Livak and Schmittgen, 2001) normalized to the average mean fold change for O4 DIV6. All measurements were performed on at least 3 cultures with each sample run in duplicate.

2.4 *Experimental design and statistical analysis*. Statistical analysis was performed using GraphPad Prism 9 (San Diego, CA). Data are presented as mean ± standard error of the mean (SEM). Variables were analyzed using one-way analysis of variance (ANOVA) with multiple analyses with Bonferroni correction for chromis-1 imaging experiments. For qPCR experiments, one-way ANOVA followed by Tukey post hoc comparisons were used to determine statistical significance. The following p-value indicate significance thresholds: *p<0.05, ** p<0.01, *** p<0.001, **** p<0.0001.

## 3. Results

### 3.1 Quantification of free zinc during oligodendrocyte development with Chromis-1 imaging

Chromis-1 imaging was used to quantify endogenous <u>free</u> zinc in oligodendrocytes from undifferentiated to fully differentiated stages. Free zinc varied depending on the developmental stage of the cells (*F*5,54 = 3.274, *p* = 0.0118). Emission ratios from developing oligodendrocytes averaged 0.59 ± 0.04 at O4-DIV2, and 0.54 ± 0.02 at O4-DIV7. An increase in endogenous free zinc was detected 1 day after cells were switched into differentiation medium on DIV6. Oligodendrocytes in differentiation media on DIV7 demonstrated a higher mean emission ratio of (0.72 ± 0.04) compared to oligodendrocytes of the same age maintained in proliferation medium (*p* = 0.044, *t* = 3.115, Figure 1A). Over the course of the following 9 days, the concentration of free zinc decreased, corresponding to increases in myelin production which begins robustly by day 13 in these cultures. Mean emission ratios decreased to 0.58 ± 0.09 at MBP-DIV9, 0.49 ± 0.04 at MBP-DIV13 and 0.52 ± 0.05 at MBP-DIV16. Free zinc was significantly lower at MBP-DIV13 compared to the peak detected at O4-DIV7 (*p* = 0.011, *t* = 3.573, Figure 1A).

**Figure 1.**
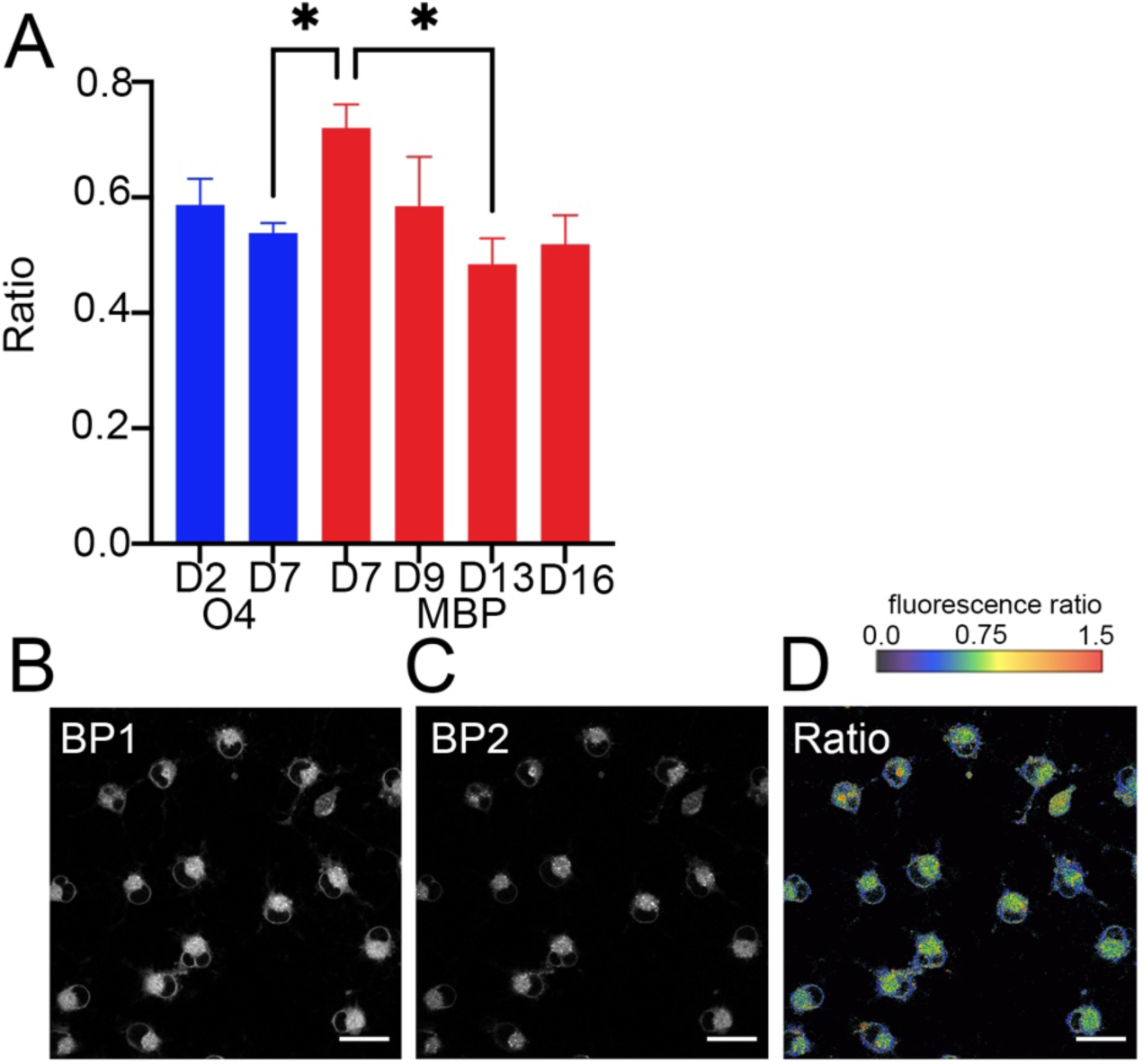
Two-photon ratiometric imaging of labile zinc reveals a transient increase following transfer from proliferation to differentiation medium. Endogenous labile zinc measurements were assessed using chromis-1 ratiometric analysis across the oligodendrocyte development lineage. (A) Summary data of chromis-1 ratio in oligodendrocytes at O4 DIV2, and DIV7 and MBP DIV7, 9, 13, and 16. Data is presented as the mean ± SEM of mean ratios per coverslip. O4 DIV2 data were collected from 168 cells from 8 total coverslips, O4 DIV7 data were collected from 741 cells from 15 total coverslips, MBP DIV7 data were collected from 329 cells from 11 total coverslips, MBP DIV9 data were collected from 159 cells from 9 total coverslips, MBP DIV13 data were collected from 575 cells from 9 total coverslips , and MBP DIV16 data were collected from 166 cells from 7 total coverslips. Chromis-1 ratios varied with developmental stage (one-way ANOVA, *F*5,54 = 3.274, *p* = 0.0118). There were significant differences between the following groups: O4 DIV7 vs. MBP DIV7 (*p* = 0.044, *t* = 3.115,) and MBP DIV7 vs. MBP DIV13 *p* = 0.011, *t* = 3.573). (B) Representative fluorescence intensity image of MBP DIV7 cells acquired with 425-562 nm (BP1) (C) 478-540 nm (BP2), and (D) ratio image (BP2/BP1). Scale bar= 20 µM.

### 3.2 Assessing developmental expression MTF-1 and MTs using qPCR

To gather additional evidence for the dynamic regulation of free zinc during oligodendrocyte development, mRNA expression was assessed for metallothioneins and the metal-responsive transcription factor (MTF-1) (Figure 2). MT1 mRNA expression was increased as oligodendrocytes differentiated (O4-DIV6: 1.28 ± 0.41, O4-DIV9: 1.36 ± 0.50; MBP-DIV9: 3.00 ± 1.24, MBP-DIV13: 8.50 ± 4.20, MBP-DIV16: 10.73±3.82; *F*(4,13) = 4.093, *p* = 0.023). MBP-DIV16 was increased compared to O4- DIV6 (*p* = 0.0357).

**Figure 2.**
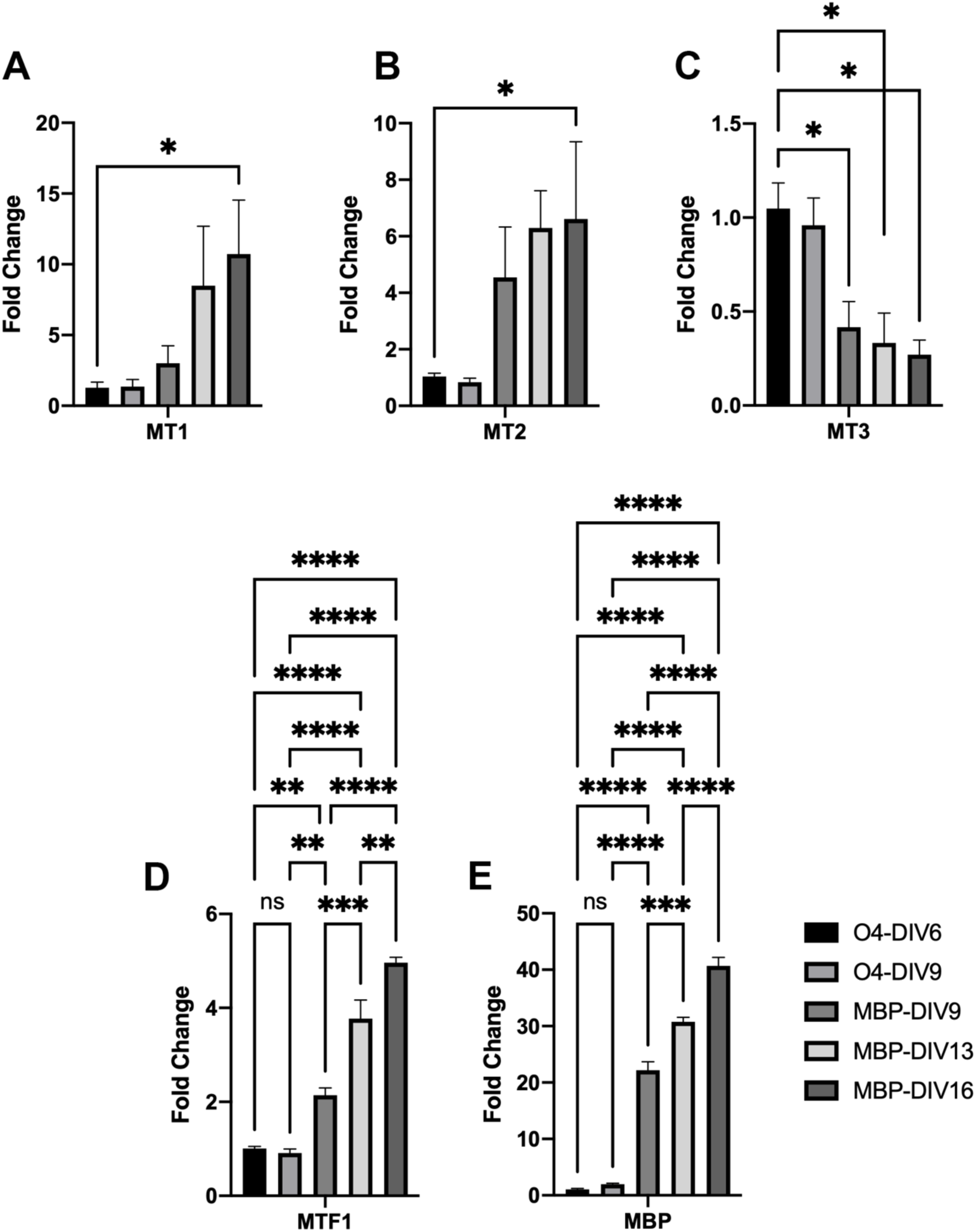
qPCR for MTs, MTF1 and MBP across OL development. Oligodendrocytes cultures were harvested 6 and 9 days following plating into medium containing PDGF and FGF (O4-DIV6, O4-DIV9) and from cultures transferred to differentiation medium containing CNTF and T3 at day in vitro 6 and harvested at 9, 13, and 16 days in vitro (MBP DIV9, MBP DIV13, MBP DIV16). qPCR was performed for MT1, MT2, MT3, MTF1, and MBP. **(A)** MT1 mRNA increased as development proceeded. **(B)** MT2 mRNA increased as development proceeded. **(C)** MT3 mRNA decreased as development proceeded. **(D)** MTF1 mRNA increased as development proceeded. **(E)** As expected, MBP mRNA increased after switch into differentiation medium. These data were collected from 3 cultures at each timepoint, except for O4-DIV6 which included 6 cultures. All analyses were performed as one-way ANOVA with Tukey’s *post hoc* analysis. *p<0.05, **p<0.01, ***p<0.001, ****p<0.0001

MT2 mRNA expression was similarly increased as oligodendrocytes differentiated O4- DIV6: 1.04 ± 0.12, O4-DIV9: 0.84 ± 0.15; MBP-DIV9: 4.54 ± 1.79, MBP-DIV13: 6.30 ± 1.32, MBP-DIV16: 6.62±2.73; *F*(4,13) = 4.920, *p* = 0.0123. MBP-DIV16 was increased compared to O4-DIV6 (*p* = 0.0391).

MT3 mRNA expression was decreased as oligodendrocytes differentiated (O4-DIV6: 1.05 ± 0.14, O4-DIV9: 0.96 ± 0.15; MBP-DIV9: 0.42 ± 0.14, MBP-DIV13: 0.33 ± 0.16, MBP-DIV16: 0.27±0.08; *F*(4,13) = 7.077, *p* = 0.0030). Compared to O4-DIV6, MT3 mRNA was decreased at MBP-DIV9 (*p* = 0.0402), MBP-DIV13 (*p* = 0.0189) and MBP- DIV16 (*p* = 0.0106).

MTF1 mRNA expression was increased very robustly as oligodendrocytes differentiated (O4-DIV6: 1.01 ± 0.05, O4-DIV9: 0.91 ± 0.090; MBP-DIV9: 2.14 ± 0.16, MBP-DIV13: 3.77 ±0.39,MBP-DIV16: 4.96±0.12; *F*(4,13) = 113.4, *p* = 0.0000). MTF-1 was increased in MBP-DIV16 compared to O4-DIV6 (*p* = 0.0000), O4-DIV9 (*p* = 0.0000), MBP-DIV9 (*p* = 0.0000) and MBP DIV13 (*p* = 0.0032). Similar increases were observed at MBP- DIV13 compared to O4-DIV6 (*p* = 0.0000), O4-DIV9 (*p* = 0.0000) and MBP-DIV9 (*p* = 0.0002). There were also increases in MTF-1 in MBP-DIV9 compared to O4-DIV6 (*p* = 0.0014) and O4-DIV9 (*p* = 0.0024).

As expected, MBP mRNA expression increased with oligodendrocyte differentiation (O4-DIV6: 1.06 ± 0.16, O4-DIV9: 1.97 ± 0.18; MBP-DIV9: 22.19 ± 1.49, MBP-DIV13: 30.76 ±0.80, MBP-DIV16: 40.70±1.50; *F*(4,13) = 464.5, *p* = 0.0000. MBP was significantly increased at all times after the switch into differentiation media. Specifically, MBP mRNA at MBP-DIV16 was higher than O4-DIV6 (*p* = 0.0000) and O4-DIV9 (*p* = 0.0000). It was also increased at MBP-DIV13 compared to O4-DIV6 (*p* = 0.0000) and O4-DIV9 (*p* = 0.0000). Even when oligodendrocytes were only in differentiation medium for 3 days for MBP-DIV9, there were increases in MBP mRNA compared to O4-DIV6 (*p* = 0.0000) and O4-DIV9 (*p* = 0.0000), indicating a rapid induction of myelination transcriptional programs.

### 3.3 Probing endogenous zinc buffer depth using the oxidizing agent DTDP

There is increasing evidence that cellular zinc levels are buffered by endogenous ligands covering a broad range of affinities (Krezel and Maret, 2007a; Colvin et al., 2010). When assessing labile zinc pools with fluorescent probes, physiologically relevant changes in total cellular zinc are thus expected to yield an attenuated response due to endogenous buffering. Because redox-active sulfhydryl-ligands play an important role in the coordination of labile zinc, (Maret, 2004) sulfhydryl-specific oxidants such as DTDP should elicit a release of zinc that parallels the depth of the labile zinc pool. To test this hypothesis, we employed chromis-1 to probe the amount of endogenously released zinc when DTDP is added to developing and mature oligodendrocytes. As illustrated with Figure 3, exposure to DTDP resulted in a large increase of the emission intensity ratio in developing O4 cells, whereas a more modest change was observed in mature MBP cells. This increase in free zinc was reversible with TPEN, except at MBP DIV16.

**Figure 3.**
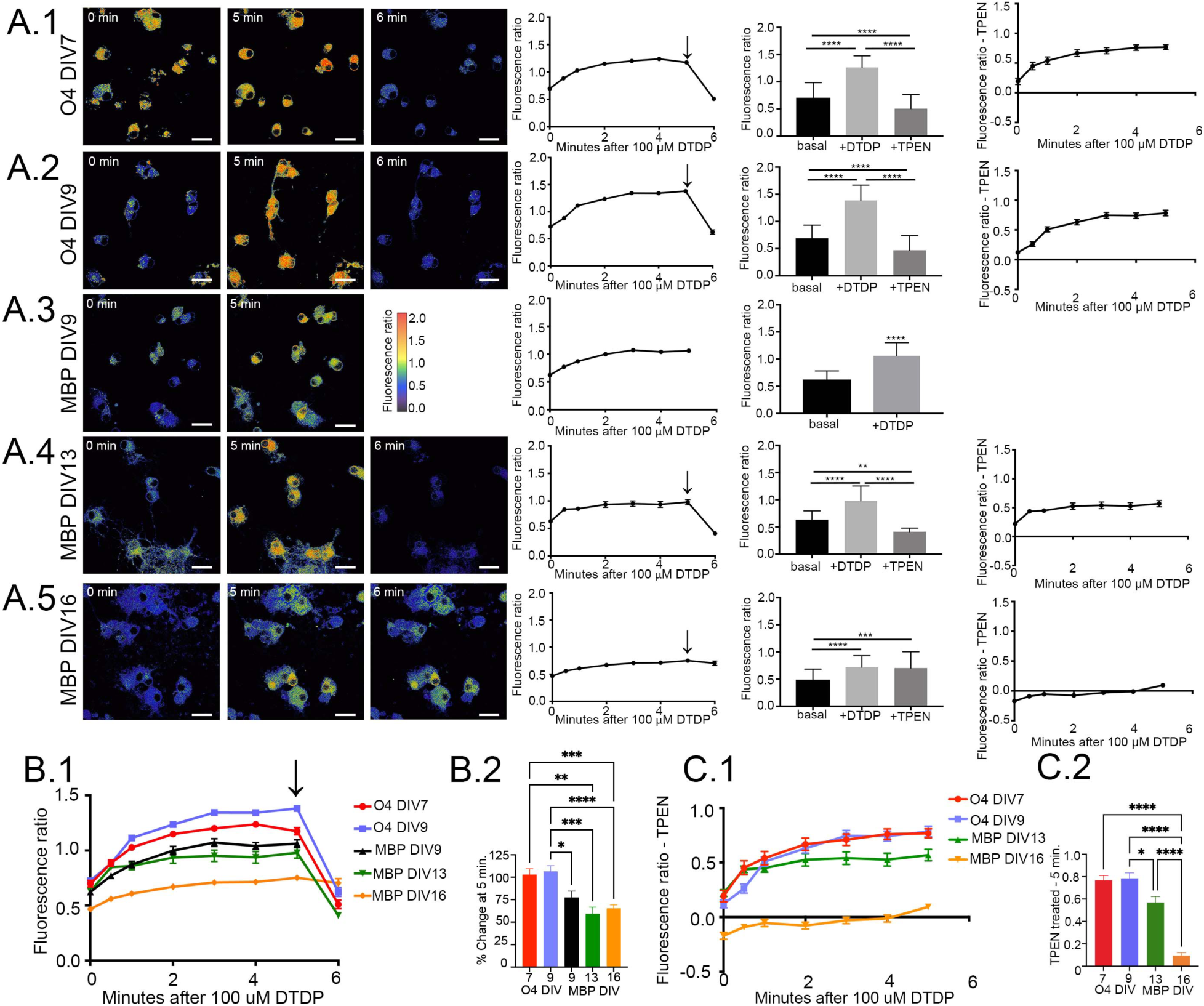
Developing oligodendrocytes release more zinc than mature oligodendrocytes when exposed to DTDP. Oligodendrocytes were exposed to 100 µM DTDP for 5 minutes, followed by 3 µM TPEN (black arrow) for one additional minute. Representative chromis-1 ratio images immediately before adding DTDP (0 min), 5 minutes after DTDP (5 min), and 1 minute after TPEN (6 min) are shown for O4 DIV7 **(A.1)**, O4 DIV9 **(A.2)**, MBP DIV9 **(A.3)**, MBP DIV13 **(A.4)**, and MBP DIV16 **(A.5)**. Data is then plotted graphically for 1) chromis-1 fluorescence ratios across time, 2) chromis-1 fluorescence ratios as bar graphs for basal (0 min), DTDP treatment (5 min) and after addition of TPEN (6 min), and 3) chromis-1 fluorescence ratios subtracting the fluorescence ratio produced by TPEN (6 min) across time. In both developing and mature oligodendrocytes DTDP produced significant increases in free zinc measured by chromis-1 fluorescence ratio. These increases were reversible by addition of TPEN, except at MBP DIV16. Bonferroni multiple comparison test, *p<0.05, **p<0.01, ***p<0.001, ****p<0.0001. **(B.1)** Summary data of chromis-1 ratio in developing and mature oligodendrocytes following exposure to 100 µM DTDP, then 3 µM TPEN (mean ± SEM) in the 5 developmental stages of oligodendrocytes. (B.2) Summary of percent change in chromis-1 ratio from 0 to 5 minutes. There were larger percent increases in the developing oligodendrocytes than mature oligodendrocytes. O4 DIV7 n= 79 cells from total 7 coverslips; O4 DIV9 n=117 cells from 9 total coverslips; MBP DIV9 n=47 cells from total 5 coverslips; MBP DIV13 n= 32 cells from 3 total coverslips; MBP DIV 16 n= 86 cells from 8 total coverslips. Bonferroni multiple comparison test, *p<0.05, **p<0.01, ***p<0.001, ****p<0.0001. **(C.1)** Summary data of chromis-1 fluorescence ratio at each timepoint minus TPEN-treated fluorescence ratio (6 minute value). **(C.2**) Summary chromis-1 fluorescence ratio 5 minutes post-DTDP treatment minus TPEN-treated chromis-1 ratio (6 min). O4 DIV7, O4 DIV9, and MBP DIV13 showed a significant decrease in fluorescence ratio following TPEN treatment that was not observed at MBP DIV16. O4 DIV7 n= 55 cells from total 4 coverslips; O4 DIV9 n= 108 cells from 8 total coverslips; MBP DIV13 n= 32 cells from 3 total coverslips; MBP DIV 16 n= 55 cells from 6 total coverslips. Bonferroni multiple comparison test, *p<0.05, **p<0.01, ***p<0.001, ****p<0.0001.

In developing oligodendrocytes in proliferation medium at DIV 7 (O4-DIV7), chromis-1 emission ratio increased 102.8 ± 6.5% following DTDP exposure, from 0.67 ± 0.03 to 1.26 ± 0.03 after 5 minutes (Figure 3A.1) (*F*7,597 = 96.5, *p* = 0.001; baseline vs 5 minutes, *p* = 0.0000, *t* = 15.4). At O4-DIV9, chromis-1 emission ratio increased 106.4 ± 6.5% from 0.73 ± 0.02 to 1.39 ± 0.02 following 5 minutes of DTDP exposure (Figure 3A.2) (*F*7,919 = 139.9, *p* = 0.0000; baseline vs 5 minutes, *p* = 0.0000, *t* = 18.5). In contrast, the increase in chromis-1 emission ratio following DTDP exposure was smaller after oligodendrocytes were switched to differentiation media. In MBP-DIV9 oligodendrocytes, the mean chromis-1 emission ratio increased 77.5 ± 7.0% following DTDP, from 0.63 ± 0.02 to 1.06 ± 0.04 after 5 minutes (Figure 3A.3) (*F*6, 317 = 26.0, *p* = 0.0000; baseline vs 5 minutes, *p* = 0.0000, *t* = 9.3). In MBP-DIV13 oligodendrocytes, chromis-1 emission ratio increased 59.2 ± 7.5%, from 0.63 ± 0.03 to 0.98 ± 0.05 after 5 minutes of DTDP exposure (Figure 3A.4) (*F*7, 248 = 23.5, *p* = 0.0000; baseline vs 5 minutes, *p* = 0.0000, *t* = 6.0). In MBP-DIV16 oligodendrocytes, chromis-1 emission ratio increased 65.4 ± 4%, from 0.47 ± 0.02 to 0.75 ± 0.02 after 5 minutes of DTDP exposure (Figure 3A.5) (*F*7, 633 = 16.6, *p* = 0.0000; baseline vs 5 minutes, (*p* = 0.0000, *t* = 8.4). Addition of the heavy metal chelator TPEN reversed the increases in chromis-1 emission ratio in O4-DIV7 cells (0.50 ± 0.04; 6 minutes vs 5 minutes, (*p* = 0.0000*, t* = 18.5), O4-DIV9 cells (0.62 ± 0.04; *p* = 0.0000*, t* = 21.0), MBP-DIV13 cells (0.41 ± 0.01; *p* = 0.0000*, t* = 9.8), but not MBP-DIV16 cells (0.70 ± 0.04; *p = NS*). MBP-DIV16 cells appear to lose their responsiveness to DTDP entirely when fluorescence related to TPEN is subtracted (Figure A5, final graph).

The percent change in chromis-1 emission ratio following five minutes of exposure to the sulfhydryl oxidant DTDP was lower in mature cells compared to developing cells (Figure 3B.1, 3B.2) (*F*4, 355 = 11.0, *p* = 0.0000). The percent increase was higher at both stages of oligodendrocytes in proliferation media compared to cells switched to differentiation media with O4-DIV7 percent change larger than MBP-DIV13 (*p* = 0.002, *t* = 3.8) and MBP-DIV16 (*p* = 0.0002, *t* = 4.3). Similarly for O4-DIV9 the percent change was also larger than MBP-DIV9 (*p* = 0.03, *t* = 3.0), MBP-DIV13 (*p* = 0.0002, *t* = 4.3) and MBP-DIV16 (*p* = 0.0000, *t* = 5.2).

Developing oligodendrocytes demonstrated a larger increase in zinc as detected by chromis-1 ratios when normalized for TPEN treatment (Figure 3C.1, 3C.2). (*F*3, 246 = 41.7, *p* = 0.0000). The ratios normalized for TPEN at O4-DIV7 (0.77 ± 0.04) and O4- DIV9 (0.78 ± 0.05) were larger than at MBP-DIV16 (0.09 ± 0.03), (*p* = 0.0000, *t* = 9.0; *p* = 0.0000, *t* = 10.6, respectively). The ratios normalized for TPEN at MBP-DIV13 (0.57 ± 0.05) were also larger at MBP-DIV16 (*p* = 0.0000, *t* = 5.4), but smaller than at O4-DIV9 (*p* = 0.0403, *t* = 2.7). Together, these chromis-1 imaging findings in response to DTDP provide evidence for a labile Zn(II) reserve in developing oligodendrocytes that decreases with maturation.

## 4. Discussion

### 4.1 Zinc increases during the transition from developing to mature oligodendrocyte

Using chromis-1 to visualize free zinc in oligodendrocytes, we demonstrated that free zinc concentrations initially increase after switching into differentiation medium, followed by a decrease in free zinc availability over time in this medium (Figure 1). Zinc is necessary for proliferation in many cell types. Recent data shows that very small fluctuations in intracellular zinc controls cell cycle regulation, with zinc deficiency inducing quiescence (Lo et al., 2020). When oligodendrocytes are in proliferation medium they continue to divide (pre-oligodendrocyte stage), but when transferred to differentiation medium proliferation ceases and myelin production begins robustly by day in vitro 13. This is also reflected in the significant upregulation of MBP mRNA (Figure 2). Oligodendrocyte progenitors are known to have an intrinsic clock with a specific number of divisions before committing to differentiation (Temple and Raff, 1986; Raff et al., 1988; Barres et al., 1994; Durand and Raff, 2000), so it possible that the zinc increase observed on transfer to differentiation medium is necessary for a final cell division. More free zinc may be required for proliferation early on in the oligodendrocyte lineage and bound zinc or sequestered zinc may be necessary to initiate later differentiation programs.

Our results may seem at odds with the increase in MTF-1 and MTs mRNA (Figure 2) which support an increase in free zinc and potentially total cellular zinc with maturation. However, chromis-1 quantifies the free or labile zinc fraction, primarily in the cytoplasm but also in certain cytoplasmic compartments such as the golgi apparatus and lysosomes (Bourassa et al., 2018). It is notably not taken up by the nucleus and there are potentially other cellular compartments, such as zincosomes (Abiria et al., 2017), which may be inaccessible to chromis-1. Furthermore, once proteins, enzymes or transcription factors bind to zinc, the zinc ions may become inaccessible to chromis-1. One explanation for the decrease in free zinc quantified by chromis-1, despite increased MT expression, is that the zinc may have entered a compartment inaccessible to chromis-1.

There are numerous compartments that might be sequestering zinc and making it inaccessible to chromis-1. The nucleus contains many zinc-finger transcription factors important in myelination programs, such as Ying-Yang-1 (He et al., 2008), myelin transcription factor 1 (Myt1) (Nielsen et al., 2004; Besold and Michel, 2015), Zfp488 (Wang et al., 2006; Soundarapandian et al., 2011), Znf161 (Sidik and Talbot, 2015), Zfp24 (Elbaz et al., 2018), and Zfp276 (Aberle et al., 2022). Nuclear zinc has also been implicated in control of epigenetic modifications (Krapivinsky et al., 2014) and these are well established to be important in controlling differentiation programs in OLs (Samudyata et al., 2020).

A second possibility is that zinc may be becoming sequestered within the myelin sheath which is deposited onto the cell culture plates of mature oligodendrocytes. Zinc is a critical structural component for myelin for maintaining myelin compaction and integrity (Inouye and Kirschner, 1984; Riccio et al., 1995; Tsang et al., 1997). Both myelin basic protein and myelin-associated glycoprotein (MAG) bind zinc which leads to stabilizing effects on MBP-membrane interactions or increased surface hydrophobicity in the case of MAG (Tsang et al., 1997; Kursula et al., 1999).

A third possibility is that zinc is binding to enzymes important in myelination programs. ERK has been shown to be important for oligodendrocyte differentiation (Ishii et al., 2012; Ishii et al., 2016) and ERK activity is regulated by a protein phosphatase that is inhibited by zinc binding (Zhang et al., 2006; Ho et al., 2008; Domercq et al., 2013). More recent work has suggested that under physiologic zinc concentrations, zinc activation of ERK is actually upstream of the phosphatase via a yet to be discovered zinc interaction with an activator protein (Anson et al., 2021).

Regardless of the specific target, it would be anticipated that activation of ERK signaling would be necessary for differentiation to proceed. Similarly, histone deacetylases (HDACs) are zinc binding proteins (Gantt et al., 2006; Kim et al., 2015) and HDACs play important roles in oligodendrocyte differentiation (Conway et al., 2012; Zhang et al., 2016). There are likely numerous other enzymes with similar zinc regulation relevant to oligodendrocyte differentiation.

### 4.2 MTF-1 and MTs increase in oligodendrocytes as they become more differentiated

The increase in MT1 and MT2 mRNA expression (Figure 2) support the hypothesis that as oligodendrocytes differentiate there is an increased mobilization of free zinc. MTF-1, the only metal-responsive transcription factor identified so far in mammals, is a cytoplasmic controller for maintaining metal homeostasis (Gunther et al., 2012). With increases in cytoplasmic zinc from either increased extracellular influx or increased intracellular release, there is increased zinc binding to the 6 zinc fingers present on MTF-1. This binding then triggers translocation to the nucleus and increased expression of the metallothioneins. The source of increased cytoplasmic zinc driving MT expression requires further study. Prior work has shown that oligodendrocyte progenitors have robust Zn^65^ uptake through a saturable influx pathway that was not blocked by calcium or glutamate antagonists (Law et al., 2003), suggesting involvement of one or more zinc transporters mediating the influx. It is possible that changes in expression or function of zinc transporters (ZnTs) or Zrt-and-Irt-like Proteins (ZIPs) underlie changes in zinc homeostasis. Available RNAseq data (see Table 1 in (Elitt et al., 2019)) suggest developmental expression of these zinc importers and exporters.

In addition to the increases in MT-1 and MT-2, there is also an upregulation of MTF-1 itself. Although there are many mechanisms of post-translational regulation of MTF-1 (Bi et al., 2006; Gunther et al., 2012), increased zinc content has been shown to enhance MTF-1 mRNA and protein expression (Hasumi et al., 2003; Chung et al., 2006; Ostrakhovitch et al., 2007). In developing myoblasts, MTF-1 expression can also be induced by increasing copper content (Tavera-Montanez et al., 2019), raising the possibility that another metal such as copper may be contributing to the MTF-1, MT-1 and MT-2 mRNA changes.

MT-3 expression has generally been confined to neurons and some subtypes of astrocytes (Fung et al., 2008), except in specific disease states where oligodendrocytes can develop inclusions positive for MT-3 (Pountney et al., 2011). MT-3 mRNA is decreasing in contrast to MT-1 mRNA and MT-2 mRNA increasing. However, MT-3 expression is not inducible by changes in metal concentrations nor driven by MTF-1 (Krezel and Maret, 2021), so it would not be expected to change based on changes in zinc or copper content. From that perspective, MT-1 and MT-2 are of greater interest in understanding metal homeostasis in oligodendrocytes. The biological implications of a decrease in MT-3 in oligodendrocytes as they differentiate requires further investigation.

### 4.3 Mature oligodendrocytes have reduced endogenous release of zinc from intracellular stores

Metallothionines are important intracellular sources of zinc and other metals and act as dynamic zinc buffers (Krezel and Maret, 2021). Based on the increase in MT1 and MT2 in mature oligodendrocytes, one might anticipate that mature oligodendrocytes would release more zinc in response to DTDP. However, our results suggest that the labile zinc pool from MTs is reduced in mature oligodendrocytes (Figure 3). One explanation is that the redox environment varies depending on the stage of oligodendrocyte development which would affect their toxicity in response to oxidative stress.

Developing oligodendrocytes are known to have reduced antioxidant capacity (Back et al., 1998; Baud et al., 2004; Folkerth et al., 2004), so it is possible that the oxidizing effect of DTDP is greater in developing oligodendrocytes. Another possibility is that zinc binding sites in MTs in mature oligodendrocytes are bound with another metal. MTs are known to have partial metalation states (Krezel and Maret, 2007b), so some sites in mature oligodendrocytes could be occupied by copper and thereby less zinc would be released in response to DTDP. More copper-rich MTs in mature oligodendrocytes would have significant toxicity implications under conditions of oxidative stress and cuproptosis is now recognized as an important form of cell death (Tsvetkov et al., 2022). New copper specific sensors and chelators (Morgan et al., 2018; Morgan et al., 2019) may help to address this question in the future. The use of zinc specific chelators combined with genetic or pharmacologic manipulation of zinc transporter or zinc storage proteins in future experiments will allow further clarification of specific zinc or copper signaling pathways in oligodendrocytes.

### 4.4 Summary

In summary, we have quantified free zinc using chromis-1 across oligodendrocyte development and demonstrated a transient free cytoplasmic zinc increase during the transition from proliferating to differentiating oligodendrocytes, suggesting zinc signals may contribute to this process. Because of the clear developmental increases in MT-1, MT-2 and MTF, maturing oligodendrocytes have an increased need for free zinc. The specific compartments, proteins, enzymes or transcription factors that sequester zinc and their relative affinities for zinc remain to be determined. Interestingly, in contrast, the depth of the endogenous zinc buffer pool in response to oxidative stress is reduced in mature oligodendrocytes which may be related to different cytoplasmic redox potential or involvement of another metal such as copper. Taken together, dynamic regulation of metal homeostasis likely drives important signaling events during oligodendrocyte differentiation.

## Acknowledgements

This work was supported in part by the National Institute of Neurological Disorders and Stroke Grant K12NS079414 (CME), the Philip R. Dodge Young Investigator Award from the Child Neurology Society (CME), internal funding from the Department of Neurology at Boston Children’s Hospital (CME), The National Institute of General Medical Science Grant R35GM136404 (CJF), the National Eye Institute Grant R01EY027881 (PAR) and the Intellectual and Developmental Disabilities Research Center at Boston Children’s Hospital (IDDRC) HD090255. Dr. Elitt serves as a consultant on the scientific advisory board of Akebia Therapeutics.

